# Direct observation of individual tubulin dimers binding to growing microtubules

**DOI:** 10.1101/418053

**Authors:** Keith J. Mickolajczyk, Elisabeth A. Geyer, Tae Kim, Luke M. Rice, William O. Hancock

## Abstract

The biochemical basis of microtubule growth has remained elusive for over thirty years despite being fundamental for both cell division and associated chemotherapy strategies. Here, we combine interferometric scattering microscopy with recombinant tubulin to monitor individual tubulins binding to and dissociating from growing microtubule tips. We make the first direct, single-molecule measurements of tubulin on- and off-rates. We detect two populations of transient dwell times, and determine via binding-interface mutants that they are separated by the formation of inter-protofilament bonds. Applying a computational model, we find that slow association kinetics with strong interactions along protofilaments best recapitulate our data, and furthermore predict plus-end tapering. Overall, we provide the most direct and complete quantification of how microtubules grow to date.

**SIGNIFICANCE:** Microtubule polymerization dynamics are fundamental to cell migration and cell division, where they are targets for chemotherapy drugs. Despite significant progress, the precise structural and biochemical events occurring at growing microtubule tips are not well defined, and better understanding is necessary for discriminating mechanisms of microtubule dynamics regulation in cells. Here, we visualize individual tubulin subunits reversibly and irreversibly interacting with dynamic microtubule tips, and thereby directly measure tubulin on- and off-rates. By analyzing plus-tip residence times of wild-type and mutant tubulin, we characterize the relative contributions of longitudinal (along protofilaments) and lateral (between protofilaments) bond energies to microtubule growth. This work provides insights into microtubule tip structure and potential modes of microtubule dynamics regulation.

## INTRODUCTION

Microtubules (MTs) are cytoskeletal polymers of αβ-tubulin that play fundamental roles in cell structure, mitosis, and intracellular trafficking(1, 2). Dynamic instability, the stochastic switching between phases of growing and shrinking, is essential for microtubule function and is directly targeted by multiple anti-cancer drugs (3–6). However, despite its fundamental importance, the kinetic basis of dynamic instability has remained elusive for more than a quarter century(7). Computational modeling(8–11), structural studies(12, 13), and reconstitution assays(14–17) have offered numerous insights, but the field still lacks a mechanistic consensus(2, 18). The missing link has been the ability to directly measure the biochemical properties of individual tubulin dimers interacting with dynamic MT tips.

The structure and composition of the MT plus-end is thought to be a critical determinant of MT assembly kinetics and catastrophe frequency. MTs are composed of tubulin subunits that form into a hollow tube of typically thirteen protofilaments(2). MT growth results from the net addition of new GTP-bound tubulin to the plus-end, where they form a GTP cap(19, 20). GTP hydrolysis in individual tubulin subunits, which is triggered by lattice incorporation, leads to changes in their structural flexibility that make the MT more prone to catastrophe (19, 21–23). Maintenance of the GTP cap is thus governed by a kinetic race between the addition of new GTP-tubulin to the tip and the loss of existing GTP-tubulin at the tip either by hydrolysis or tubulin dissociation. However, the fundamental rates at which GTP-tubulin subunits associate with and dissociate from MT tips has remained unknown because the structurally heterogeneous growing MT tips are biochemically complex, and because it has not been possible to directly observe with high temporal resolution individual tubulins interacting with the growing tip.

The structural and biochemical challenges posed by growing MT tips can be understood as follows. By “tip”, we refer to the thirteen tubulin dimers at the plus-end terminus of each protofilament in the polymer. Some protofilaments extend farther than others (24), so tubulins on the tip can be stabilized to different degrees depending on the number of intra- and inter-protofilament contacts they make(14). A further layer of complexity comes from the fact that the tubulin subunits themselves undergo conformational changes during and after their addition to the MT, and these changes modulate their biochemical properties in unknown ways (2). This structural and biochemical heterogeneity at the tip makes it difficult to extrapolate microscopic states and transient events from ensemble kinetic measurements and static structures (14, 25–27). The result is that a number of plausible models for microtubule growth have been proposed that are based on different and sometimes conflicting assumptions(8, 12, 14, 18, 22, 28). Single-molecule measurements have the potential to reveal kinetic heterogeneity in complex systems (29–31), but numerous technical difficulties have to date prevented the extension of these methods to tubulin and microtubules.

In the current work, we overcome the longstanding challenge of single-molecule tubulin measurements by applying two recently-developed technologies—recombinant tubulin and interferometric scattering microscopy (iSCAT). We first create biochemically homogeneous populations of tubulin subunits sparsely labeled with 20-nm gold nanoparticles. We then visualize dynamic MTs label-free using iSCAT, and watch in real-time at high frame-rates as the gold-labeled subunits transiently bind to the tips. In this way we directly quantify the rates of tubulin association and dissociation. Interestingly, the transient dwell times of tip-bound tubulin split into two sub-populations, indicating that our measurements are reporting on two classes of binding sites at the MT tip. Experiments using mutant tubulin demonstrate that perturbing the inter-protofilament contacts reduces the fraction of long duration events. We thus interpret our fast and slow events as those having zero and one lateral contacts, respectively. Applying a computational kinetic model, we arrive at a self-consistent parameterization of MT growth that supports the formation of ragged and tapered tips. Overall, our results provide the first viable approach for quantifying the long-elusive tubulin-microtubule interactions that underly MT growth and dynamic instability.

## RESULTS

### Monitoring individual tubulins within growing microtubules with iSCAT microscopy

We combined two recently introduced technologies, recombinant tubulin(32) and interferometric scattering microscopy (iSCAT)(33–36), to make single-molecule observations of individual tubulin dimers binding to growing MT tips. We first generated a homogenous population of *S. cerevisiae* tubulin biotinylated through a C-terminal KCK epitope (tubulin-KCK) on the β-subunit (Tub2) (Fig. 1A). We then mixed with 20-nm streptavidin-coated gold nanoparticles (1:2000 gold:tubulin ratio), and measured microtubule dynamics *in vitro* (Fig. 1B). MTs were detected label-free, as expected(35–37), and tubulin-bound gold nanoparticles (tubulin-gold) appeared as points of high contrast (Fig. 1C). To isolate the growth reaction and avoid more complex behaviors associated with catastrophe, we performed our assays using GTPγS, the hydrolysis-resistant GTP analog that best supports growth and prevents catastrophe for yeast tubulin(38, 39). Individual tubulin-gold landing events at MT tips were clearly observable (Fig. 1C, Movie S1). These landings reflected end-binding events, because no landings were observed along the body of the microtubule. Thus, our labeling and iSCAT approaches provide a viable way to study MT assembly kinetics at the single-molecule level.

**Figure 1.**
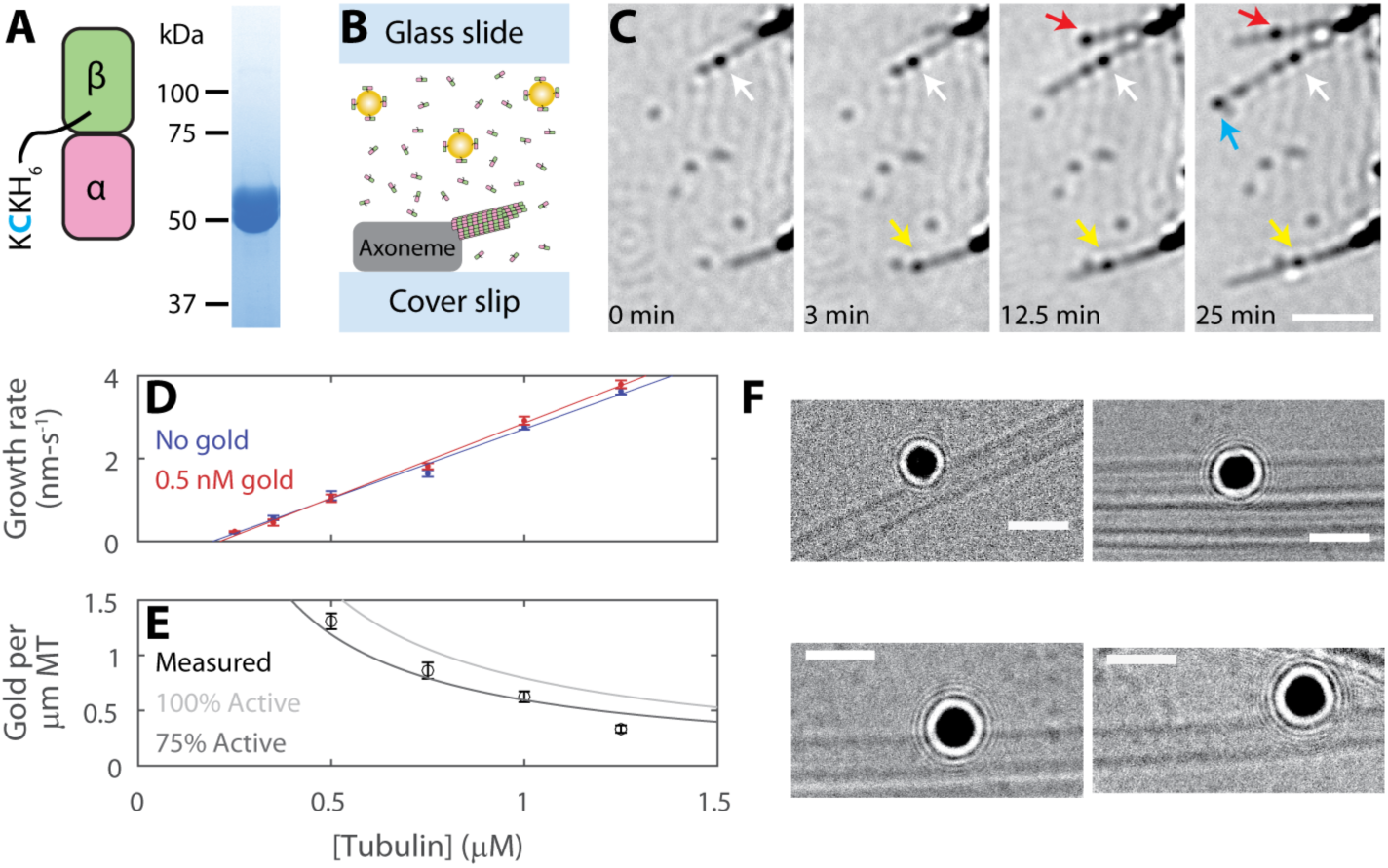
Gold-nanoparticle-labeled recombinant tubulin incorporates into microtubules without perturbing kinetics or structure. (**A**) Recombinant yeast tubulin was prepared with a biotinylated C-terminal KCK extension. Gel shows Tubulin-KCK purity. (**B**) Diagram of the *in vitro* reconstitution of MT growth. Sea urchin axonemes seed polymerization; microtubules formed from yeast tubulin sparsely tagged with 20-nm gold nanoparticles. (**C**) Time lapse iSCAT images of growing MTs and tubulin-gold association. Arrows indicate tubulin-gold incorporation events. MTs readily grow past incorporated tubulin-gold. Axonemes appear as dark structures on the right of each frame. Scale bar 2.5 μm. (**D**) MT growth rates for tubulin-KCK with and without 0.5 nM of gold nanoparticles. Data shown as mean ± standard error of the mean (SEM) for N=6-27 growing MTs. Solid lines show linear fits to data. (**E**) The number of gold nanoparticles per micron of polymerized MT. All experiments performed with a constant 0.5 nM of gold. Data shown as mean ± SEM for N=31-61 MTs. Lines show the expected number of gold per micron of microtubule given the tublin:gold ratio and assuming 8 nm per tubulin and 13 protofilaments per microtubule. Percent active refers to the portion of the gold nanoparticle pool that is not lost to nonspecific surface binding. (**F**) Example cryo-EM micrographs showing that gold-labeled tubulin can incorporate into the wall of microtubule without causing observable defects. Scale bars 50 nm.

Multiple lines of evidence indicate that the gold-labeling did not substantially perturb the kinetic or structural properties of the labeled tubulins as they polymerize into MTs. First, the presence of gold did not alter microtubule growth rates, indicating that tip-bound tubulin-gold did not alter the binding kinetics for subsequent unlabeled tubulin-KCKs (Fig. 1D). Second, tubulin-gold incorporated into the MT lattice at a spatial frequency that closely corresponded to their population as a fraction of total tubulin (Fig. 1E). Thus, tubulin-gold is not disadvantaged relative to tubulin-KCK for incorporating into growing MTs. Finally, cryo-EM imaging of MTs with tubulin-gold polymerized into the wall revealed normal lattices that were free of obvious defects (Fig 1F, Fig S1).

### Direct observation of the tubulin on-rate constant

We analyzed individual tubulin-gold binding events as they occurred in real time by recording microtubule growth at 100 frames per second. We also imaged at 1,000 frames per second, but these data were not useable because the frequency of false positive events arising from the relatively slow diffusion of gold nanoparticles was too high (Fig. S2). We observed two classes of binding events: reversible ones where tubulin-gold transiently dwelled at the MT tip before dissociating (Fig. 2A and Movie S2), and irreversible ones where the tubulin-gold was integrated into the MT lattice (Fig. 2A and Movie S3). Counting the raw number of irreversible tubulin-gold binding events per tip per second at different total tubulin concentrations allowed us to predict the unlabeled MT growth rate to within a factor of about two, further validating our single-molecule approach (Fig. S3). Next, we scored the fraction of observed events that were irreversible at each tubulin concentration (Fig. 2B). This irreversible fraction increased with increasing tubulin concentrations, reflecting a higher likelihood of ‘trapping’ by incoming subunits. Finally, we estimated the on-rate constant for tubulin association by counting the total number of binding events (reversible and irreversible) per tip per second and dividing that number by the gold concentration of 1 nM (Fig. 2C). The mean on-rate constant was 3.4±1.6 μM^-1^s^-1^ per tip or 0.26 μM^-1^s^-1^ per protofilament (Fig 2C, fit ± 95% confidence intervals), and was independent of the total tubulin concentration, as expected. To our knowledge, this is the first direct measurement of the tubulin on-rate constant, and is the only measurement that does not require fitting to a model(14).

**Figure 2.**
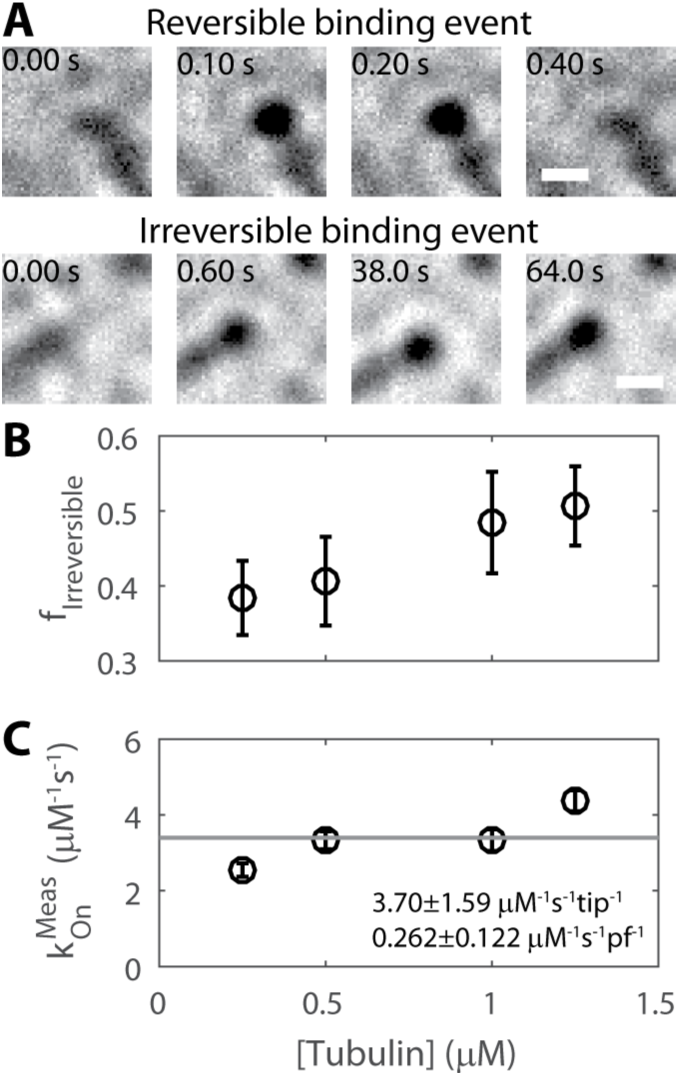
Visualization of single tubulin-gold binding events under iSCAT at 100 frames per second. (**A**) Example images showing reversible and irreversible tubulin-gold binding events. (**B**) The fraction of irreversible events observed at each free tubulin concentration. Data shown as mean ± SEM for N=22-34 flow cells. (**C**) Direct measurement of the tubulin-gold on-rate constant, determined by counting the total number of tubulin-gold binding events per second per μM gold. Data shown as mean ± SEM for N=22-34 flow cells. Value converted to per protofilament (pf) basis by dividing by 13. All measurements with 1 nM of gold.

### Direct observation of the tubulin off-rate reveals two populations

We next characterized the tubulin off-rate from the tip by quantifying the dwell times of the reversible tubulin-gold binding events at four different tubulin-KCK concentrations (Fig. 3A-B). Three trends were apparent from this analysis (Fig. 3B): (1) Dwell times spanned four orders of magnitude from 10^-2^ s to 10^2^ s (2) No distribution followed a single-exponential, and (3) Increasing concentrations of tubulin-KCK led to shorter dwell times. To interpret the distribution of measured dwell times, F(τ), the data were fit to a biexponential (Fig. 3C):

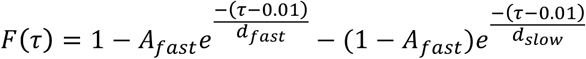

where A_fast_ is the proportion of fast-phase events, d_fast_ is the fast-phase dwell time, d_slow_ is the slow-phase dwell time, and the x-offset of 0.01 accounts for the frame rate (residuals in Fig. S4). Both the fast and slow durations decreased substantially with increasing tubulin concentration (Fig. 3D and Fig. S5). The loss of long dwell times at high tubulin concentrations is consistent with the corresponding increase in the fraction of irreversible binding events noted above (Fig. 2B).

**Figure 3.**
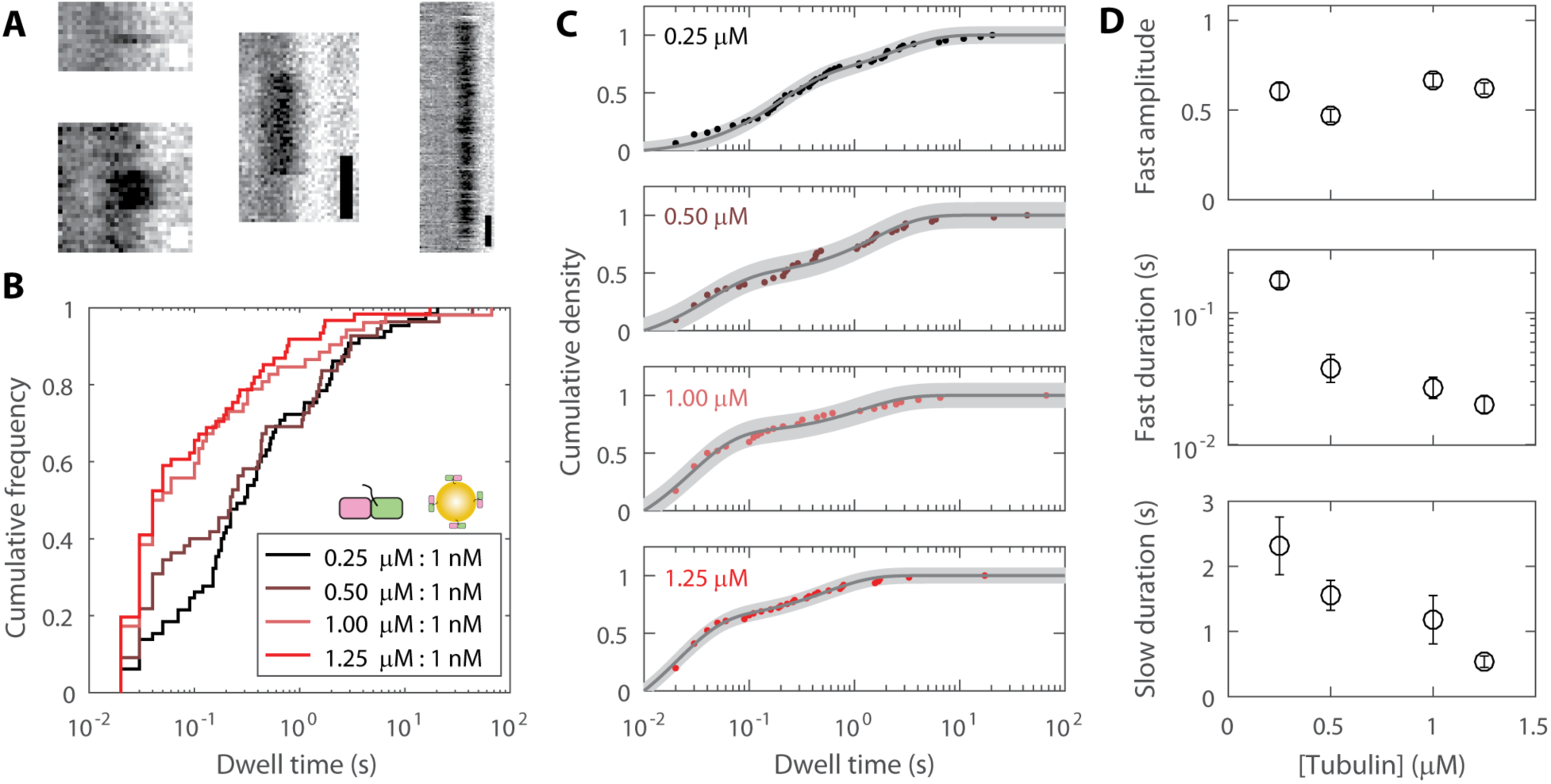
Quantification of transient tubulin-gold tip-dwell times. (**A**) Example kymographs showing reversible tip-dwell events. White scale bar is 50 ms and black scale bar is 200 ms. (**B**) Distributions showing all reversible dwell time data for four total tubulin concentrations, all with 1 nM of 20-nm gold nanoparticles (N=52-66 dwell events). (**C**) The data were well-fit with biexponentials. Uncertainty in the fits are shown as shaded gray areas. (**D**) Best parameters determined from the biexponential fit, shown as fit value ± 95% confidence intervals (CI).

### Analysis of mutant tubulins with perturbed binding interfaces reveals that inter-protofilament bonds are necessary for long dwell times at microtubule tips

The presence of two dwell time populations suggests that there are (at least) two distinct classes of tubulin-binding sites on the tip. But what intra- or inter-protofilament interactions define these classes? To address this question, we took advantage of tubulin mutants with impaired longitudinal or lateral interfaces (Fig 4A): ΔLong-KCK has two mutations at its plus-end that attenuate longitudinal tubulin-tubulin interactions(32), and ΔLat-KCK has a single mutation that weakens the lateral binding interface(40). We measured microtubule growth at multiple concentrations of wild-type tubulin (no KCK tag, not biotinylated) in the presence of 10 nM of biotinylated mutant tubulin and 1 nM of 20-nm streptavidin-coated gold nanoparticles (Fig 4A). Adding gold-labeled mutant tubulin modestly decreased the MT growth rate compared to the control (Fig 4B); however, it strongly decreased the number of gold counted per micron of MT polymer (Fig. 4C). Hence, the mutants tended to transiently poison MT tips, with the longitudinal interface mutant yielding a stronger effect. We quantified reversible dwell times for each mutant, focusing on the highest tubulin concentration (1.25 μM) where mutant incorporation was highest. The ΔLong-KCK dwell time distribution was not significantly different than that of 1.25 μM tubulin-KCK (Fig. 4D-E), as expected since none of the tip-binding interfaces were mutated. This result rules out the possibility that slow-phase dwells were due to transient burying of the labeled tubulin by unlabeled tubulin. The second mutant, tubulin-ΔLat-KCK, had identical fast and slow dwell times as tubulin-KCK, but the fast amplitude increased by about 50% (Fig. 4D-E and S6). This shift in the balance of fast and slow dwells indicates that half of the slow duration events were converted to fast duration events. Thus, slow and fast phase dwells differ due to the addition of one lateral bond (Fig. 4E).

**Figure 4.**
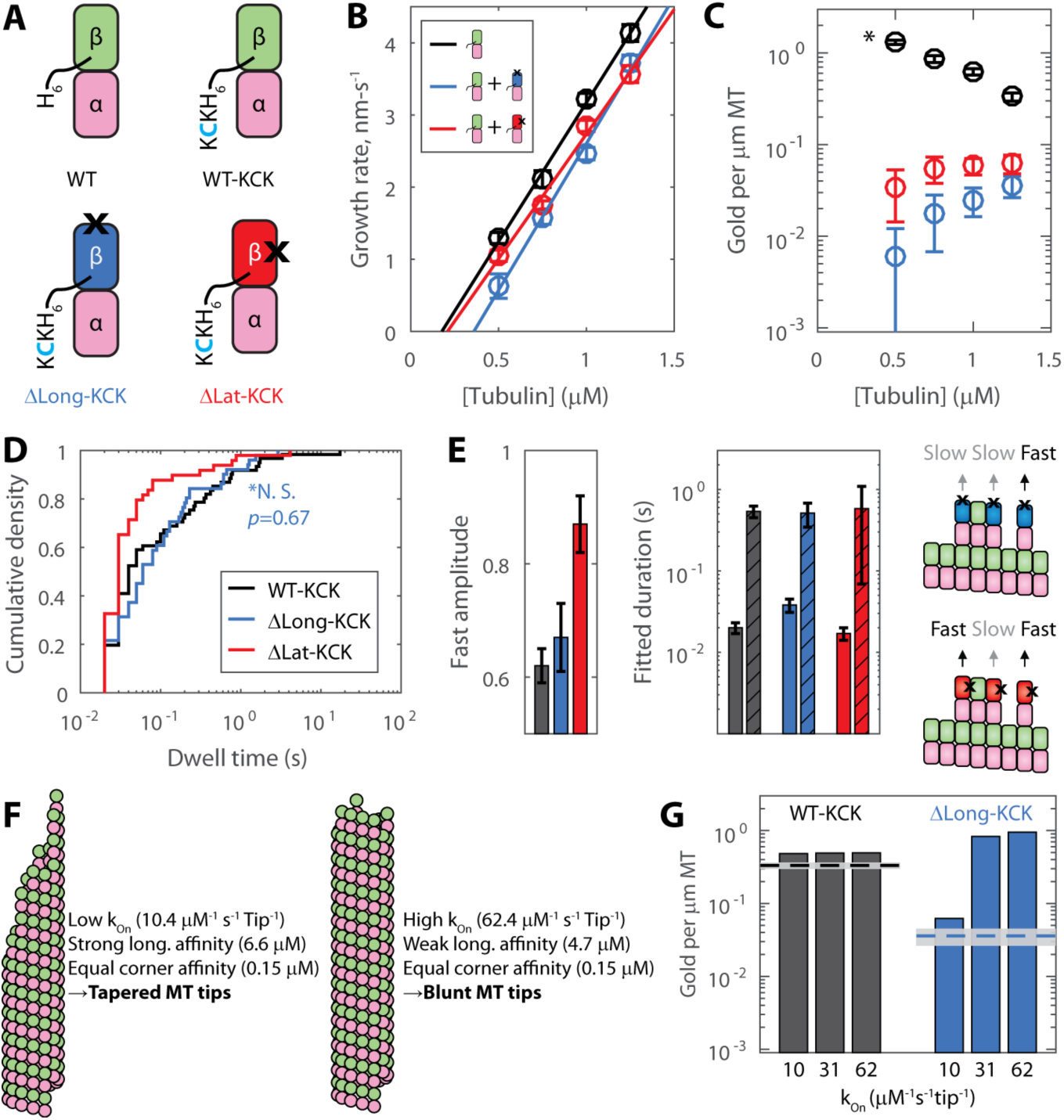
Experiments and computational modeling with mutant tubulin support tapered tip growth. (**A**) Library of tubulin species. Only those with a C-terminal KCK can bind to gold nanoparticles. Black X indicates the locations of interface-perturbing mutations. (**B**) Observed growth rate as a function of free wild-type tubulin with or without 10 nM mutant tubulin and 1 nM 20-nm gold nanoparticles. Data shown as mean ± SEM for N=20-29 MTs for wild-type alone, N=9-24 for ΔLong-KCK added, and N=17-37 for ΔLat-KCK added. (**C**) The number of gold per micron of MT for tubulin-KCK (mean ± SEM, N=31, 35, 61, and 47 microtubules measured for 0.50, 0.75, 1.00, and 1.25 μM tubulin, respectively), ΔLong-KCK (N=45, 88, 98, 92), and ΔLat-KCK (N=50, 65, 77, 63). The asterisk indicates that two-fold more gold was used in the mutant experiments versus the wild-type. (**D**) Tip dwell times of gold-labeled ΔLong-KCK (blue, N=51) and ΔLat-KCK (red, N=49) on wild-type MT tips versus control (tubulin-KCK, black N=61) at 1.25 μM total tubulin. A Wilcoxon rank sum test shows the tubulin-KCK and ΔLong-KCK distributions are not significantly different. (**E**) Results of biexponential fits to wildtype and mutant dwell time data (fits ± 95% CI, colors match legend in (D)). Diagonal lines denote slow phase dwell times. Diagram to right rationalizes why the fast phase amplitude increased about 50% for ΔLat-KCK. (**F**) In our computational model, different assumed k_On_ values yielded vastly different longitudinal affinities, leading to different predicted tip structures. In the cartoon MT ends, the variation in protofilament extension length matches that observed in the respective simulations. (**G**) Predicted (bars) and experimental (lines, mean ± SEM) number of gold per micron at 1.25 μM tubulin. Three different on-rate regimes for the computational model shown (includes a 0.75× correction factor for percent active (Fig. 1E)).

### A computational kinetic model supports a slow tubulin on-rate constant

To obtain insight into the biochemical mechanisms underlying our observations, we introduced a minimal computational kinetic model of microtubule elongation that includes a tubulin on-rate constant (k_On_) and two adjustable parameters that describe the strength of longitudinal and lateral contacts (41, 42). For each simulated k_On_, the strength of the two interfaces were refined by global fitting to the experimental growth rates and dwell times (see Fig. S7 and methods). A range of k_On_ values yielded good fits that had similar ‘corner’ affinities (simultaneous longitudinal and lateral interactions) but that varied in the relative strength of longitudinal and lateral interactions. Smaller k_On_ values (closer to our experimentally measured value; Fig. 2C) required a stronger longitudinal affinity (corresponding to a slower off-rate), and resulted in rough and tapered MT end configurations (Fig. 4F). In contrast, faster k_On_ values required a weak longitudinal affinity (corresponding to a fast off-rate), and resulted in blunt MT end configurations where growth is dominated by addition at ‘corner’ sites (Fig. 4F and S7).

To choose between these contrasting scenarios, we extended our simulations to model our mutant tubulin experiments (Fig. 4A). We assumed that mutants interacted with the MT identically to wild-type, except on the perturbed interface, where we imposed a penalty to attenuate the lateral or longitudinal interaction strength. We then sought to determine the attenuation factor for each mutant by fitting to the mutant growth rate data (Fig. S8). For the longitudinal mutant simulations, we obtained an attenuation factor on the order of 10^6^-10^7^. This attenuation is consistent with the observation that the double mutant cannot assemble MTs by itself (32), and is plausible given that single mutations are commonly observed to decrease binding affinity by three or more orders of magnitude(43). Even with attenuation factors that completely cancel the lateral interface contribution to binding affinity, we were unable to recapitulate the decrease in growth rates observed for that mutant (Fig S8). Consequently, we focused on the simulations with the longitudinal mutant. We used our simulations to predict the expected number of gold per micron MT, and found that only low k_On_ parameterizations were capable of replicating the experimental data (Fig. 4G and S9). Hence, the parameterization that best described our entire data set was k_On_=10.4 μM^-1^s^-1^tip^-1^, K_D,Long_=6.61 μM and K_D,corner_=0.147 μM, consistent with ragged MT tip tapers during growth (Fig. 4F). We note that this upper limit value for k_On_ is approximately three-fold higher than our experimentally measured value, but view this as a good match because numerous experimental factors, including loss of gold to non-specific surface binding and tubulin degradation over time, support the measured value as a lower limit.

## DISCUSSION

The current work presents the first direct, single-molecule interrogation of the biochemical mechanisms that dictate microtubule growth kinetics. Our determinations of the tubulin on-rate constant (Fig. 2C) and off-rate (Fig. 3) were made by observing the fates of individual tubulins as they transiently interacted with the microtubule tip, and hence required no model fitting or assumptions. Our off-rate measurements provide insight into the complex biochemistry of the growing microtubule tip: Two dwell-time populations were observed, consistent with tubulin reversibly associating through either a single longitudinal interaction or at a ‘corner’ site, and these structural interpretations were supported both by results from mutant tubulin and by kinetic simulations.

Our kinetic measurements provide constraints for the energetics underlying the stability of the GTP cap at dynamic MT tips. While it is established both lateral and longitudinal tubulin-tubulin interactions contribute to the stability of the microtubule lattice, the relative contributions of lateral and longitudinal free energy are debated(2). Based strictly on the experimental on-rate and off-rate measurements, the data predict a longitudinal bond energy of 10.9 k_B_T and a lateral bond strength of 2.5 k_B_T, values which lie in the middle of various published models(8–10, 14, 18). Our values suggest that the longitudinal bond energy is strong enough to be a major contributor to MT elongation, but not strong enough to drive the elongation of individual protofilaments or unseeded nuclei at modest tubulin concentrations. Our measurements do not, however, inform on the curvature or mechanical states of the protofilaments at the tip, necessitating further studies.

A notable result from our kinetic simulations in the identified slow k_On_ regime is the natural occurrence of tapered tip structures during growth. Tapered end configurations that have been observed by electron microscopy (18, 24, 44–47), inferred from fitting the intensity profile of fluorescently-labeled MTs (14, 48), and observed indirectly in the form of EB comet splitting in the presence of eribulin(49). Our data now outline the biochemical mechanisms by which these tapers grow. Key to taper formation is the ‘associative trapping’ of tubulin subunits, described as follows. The longitudinal affinity of tubulin is sufficiently strong such that a newly-bound subunits with no lateral contacts can stay bound for a few tenths of a second – long enough that there is a low but significant probability that a second tubulin will bind in at one the two open lateral positions. The occasional rare event where such lateral-stabilization occurs leads to longer-lived binding (over a second), and can thereby initiate a new taper. Associative trapping occurs with increasing frequency at higher tubulin concentrations, underlying the increase in the irreversible fraction (Fig. 2B) and loss of long-lived dwell events (Fig. 3D) seen in the experimental data. A low k_On_ model with associative trapping contrasts high k_On_ models, which suggest that only tubulins that land in corner configurations persist long enough to contribute to growth. We note that at the identified k_On_ regime, very few (about 3%) transient interactions at the tip were missed at our chosen frame rate, reinforcing the idea that iSCAT is able to capture the transient tubulin interactions that drive MT growth.

The fundamental tubulin on- and off-rates that we identified in this study form the basis for GTP cap maintenance during microtubule growth. The on-rate measured here sets an upper-limit for the rate at which GTP hydrolysis could be occurring without depleting the cap, but further experiments are necessary to gain a full understanding of how these events coordinate to drive dynamic instability. Microtubule polymerization mechanisms are conserved from yeast to humans, with specific kinetic rates varying with species, isoform, and post-translational modifications(2, 50, 51). Thus, the single-molecule approach developed here holds promise for discovering the biochemical origin of these differences, as well as for defining the mechanisms by which various polymerases, depolymerases, and chemotherapeutic drugs exert their effects by altering tubulin on- and off-rates at the MT plus-end.

## MATERIALS AND METHODS

### Protein preparation

Plasmids to express wild-type yeast αβ-tubulin were previously described(32, 38, 52). Plasmids to express wild-type yeast αβ-tubulin with a C-terminal ‘KCK’ tag, a β:F281A mutation of Tub2p (yeast β-tubulin) with C-terminal ‘KCK’ tag (main text ΔLong-KCK), and a β: T175R, V179R mutation of Tub2p (yeast β-tubulin) with C-terminal ‘KCK’ tag (main text ΔLatX-KCK) were made by QuikChange (Stratagene) mutagenesis, using the expression plasmid for wild-type Tub2 as template and with primers designed according to the manufacturer’s instructions. The integrity of all expression constructs was confirmed by DNA sequencing. All wild-type and mutant yeast αβ-tubulins (wild-type-KCK, β:T175R/V179R-KCK, β:F281A-KCK) were purified from inducibly overexpressing strains of *S. cerevisiae* using Ni-affinity and ion exchange chromatography, as previously described(32, 38, 41, 52). After purification, all ‘KCK’ samples were labeled with EZ-Link Maleimide-PEG2-Biotin (Thermo Scientific). All tubulin samples were incubated with 2 mM TCEP for 30 min on ice. Following incubation with TCEP, 20-fold excess of Maleimide-PEG2-Biotin was added and samples were incubated for 2 hours at 4°C. Samples were exchanged into storage buffer (10 mM PIPES pH 6.9, 1 mM MgCl_2_, 1 mM EGTA) containing 50 μM GTP with 2 mL, 7K MWCO Zeba spin desalting columns (Thermo Scientific). Tubulin samples were then flash-frozen and stored at -80°C. The degree of tubulin subunit biotinylation was determined to be within error of 100% (one biotin per tubulin dimer) using a fluorescence biotin quantification kit (Thermo-Fisher 46610). Prior to any assay, tubulin was thawed, spin-filtered (100 nm pore size) to remove aggregates, and the concentration was re-measured by absorbance at 280 nm.

### Interferometric scattering microscopy

An iSCAT microscope was custom-built for this study. The design roughly followed the scanning illumination design suggested by the Kukura Lab(33, 34), and the system was built around an existing total internal reflection microscope(53). Illumination was provided by a 650 nm laser (OdicForce OFL384), which was mode cleaned using a single-mode optical fiber (Thorlabs). Scanning was performed using an acousto-optic deflector (Gooch and Housego 45070-5-6.5DEG-.633). A 60× 1.49 numerical aperture objective (Nikon) equipped with an objective heater (Bioscience Tools) was used. Images were taken using a Basler Ace CMOS camera (AcA2000_165um) accessed by custom-written LabVIEW software. The total microscope magnification was 167x, resulting in an image calibration of 33 nm/pixel. For all experiments performed, unless otherwise stated, the camera exposure time was 9.435 ms, the laser was scanned over the sample 290 times per exposure, and the power incident on the sample was ∼25 W/cm^2^. All images and movies shown in this study were flat-fielded(33, 34), and thus pixel intensity represents a percent contrast rather than a raw grayscale count.

### Microtubule growth and reversible tubulin binding assays

Microtubule growth assays under iSCAT were based off of established protocols(38). MTs were polymerized in flow chambers off of sea urchin axoneme seeds. Cover slips were washed with ethanol and silanized using 1H,1H,2H,2H-Perfluorodecyltrichlorosilane (Alfa-Aesar) before being constructed into ∼10 μL flow cells using double-sided tape. Chambers were first washed with 20 μL of BRB80 buffer (80 mM K-PIPES, 1 mM EGTA, 1 mM MgCl_2_, pH 6.8), and then axoneme seeds were added. After five minutes, a blocking buffer (5 mg/mL BSA and 0.25 mM GTPγS in BRB80) was added. After ten minutes, polymerization assay mix was added and the flow chamber was sealed using clear nail polish. The polymerization assay mix included a controlled amount of biotinylated tubulin, 2 mg/mL BSA, 2 mM GTPγS, 0.4 mM MgSO_4_ and 0.08% methylcellulose in PEM buffer (100 mM PIPES, 1 mM EGTA, 1 mM MgSO_4_, pH 6.9). Streptavidin-coated 20-nm gold nanoparticles (BBI Solutions) were added to the polymerization assay mix at a final concentration of either 0.5 nM or 1.0 nM. In all cases, streptavidin-coated gold and biotinylated tubulin were pre-mixed on ice for at least 10 minutes. In all assays, a large stoichiometric excess of tubulin was used (ranging from 10:1 to 2,500:1 tubulins per gold). In experiments with biotinylated mutant tubulin, mutants were pre-mixed with gold at a 10:1 ratio on ice for 10 minutes before unbiotinylated wildtype tubulin was added to the polymerization mix (final concentration of 0.25-1.25 μM wild-type tubulin, 10 nM biotinylated mutant tubulin, 1 nM 20-nm streptavidin gold). Flow chambers were heated to 30°C in a mini incubator (Bio-Rad), and loaded onto the iSCAT microscope, where temperature was maintained using an objective heater. Sample temperature was measured periodically using a digital thermometer (Leaton L812) to ensure consistency. Growth rate movies were acquired at 2 frames per second. To assess gold per micron microtubule, 20 frame stacks were taken at 100 frames per second (microtubule fluctuations are visible at that frame rate, and aid in distinguishing whether a gold is truly on a microtubule, or just has stuck to the surface near a microtubule). Growth rates and the number of gold per micron microtubule were measured in ImageJ. For all experiments performed, data were taken from at least two flow cells spread over at least two separate experimental days.

To measure individual tubulin binding and unbinding events, the tubulin growth assay was repeated with a controlled amount of tubulin-KCK and 1 nM of streptavidin coated 20-nm gold nanoparticles (BBI Solutions). For the mutant tubulin experiments, the 1 nM of gold was covered with 10 nM of biotinylated mutant tubulin, and the rest of the tubulin in the pool was nonbiotinylated. Experimentally, a growing tip was located, and a 20,000-frame movie was taken at 100 frames per second. Only axonemes containing clear, single growing microtubules (determined by contrast and fluctuations) were recorded. Multiple movies of multiple tips were taken in each flow cell, and no flow cell was imaged for long than ∼40 minutes to minimize tubulin loss (to nonspecific surface binding, aggregation, or denaturation). Movies were fed into ImageJ and kymograph analysis was performed manually. Most movies had 0, 1, or 2 events. An event was labeled as irreversible if the microtubule was seen to grow past the tubulin-gold, or if the dwell event persisted throughout the lifetime of the flow cell (significantly longer than 200 seconds). Only events that were at least two frames long were scored.

### Cryo electron microscopy

To view the overall MT architecture in the presence of gold label, biotinylated tubulin was polymerized in 1× G-PEM (100 mM Pipes, 2 mM EGTA, 1 mM MgCl_2_, 5% glycerol, pH 6.9) with 1 mM GTPγS and 1 nM of 20-nm streptavidin gold nanoparticles at 30°C for 2 hr. The tubulin to gold ratio was 1000:1. Cryo-EM grids were prepared by applying 3 μL of MTs to a glow-discharged Quantifoil R1.2/1.3 300-mesh gold holey carbon grid (Ted Pella Inc.) and blotted for 4.0 s under 100% humidity at 30°C before being plunged into liquid ethane using a Mark IV Vitrobot (FEI). Micrographs were collected on a Talos Arctica microscope (FEI) operated at 200 kV with a K2 Summit direct electron detector (Gatan). Serial EM software (David Mastronarde, J. Richard McIntosh, http://bio3d.colorado.edu/SerialEM/) was used to manually collect data. Images were recorded at 6,700X and 13,500X magnification using a defocus from -2.5 μm to -3 μm. All micrographs were dose-fractionated to 20 frames with a dose rate of about 6 electrons per pixel per second, with a total exposure time of 10 s. Micrographs were processed using RELION v2.1(54): images were motion corrected using MotionCor2 and CTF estimation was performed using CTFFIND4.

### Computational modeling

We created a computer program to perform kinetic Monte Carlo simulations of growing MT plus ends. The model is similar to one we used previously(41) and to an earlier implementation(10). Briefly, the MT lattice is represented by a two dimensional array with a periodic boundary condition to mimic the cylindrical wall of the MT. MT elongation is simulated one biochemical reaction (subunit association or dissociation; GTPase is not considered in this model because our experiments used a hydrolysis-resistant GTP analog) at a time. The association can happen at the tip of each protofilament, and association rate is given by k_On_ × [tubulin]. The occupied terminal subunits can dissociate from the MT lattice at a rate given by k_on_ × K_D_, where K_D_ is the affinity determined by the sum of all tubulin-tubulin interactions as described previously(10, 41, 42). After assuming a value for k_On_, the strengths of the longitudinal interaction were determined by fitting the model predictions on growth rates and dwell times to the experimental values. The goodness of the fit is determined using a weighed sum of the squared residuals. The weight of growth rate data points are given by 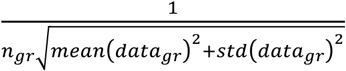, where n is the number of growth rate data points. The weight of dwell time data points are given by 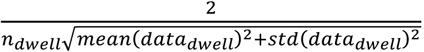. The best fitting parameter set was found using the Gauss-Newton Algorithm to minimize the weighted sum of the residuals. To ensure that the minima found is the global minimum, we used at least 25 different initial parameter sets.

To simulate growth rates in the presence of interface-perturbed mutants, we modified the simulation code to include an additional species of tubulin with a user-specified concentration. We assumed that the interface mutants decrease the strength of interaction (affinity) at that interface. This attenuation affects the dissociation rate of the affected subunit, but not its association rate. For ΔLong-tubulin (β: T175R, V179R mutations), longitudinal interactions involving the plus-end of the mutant are attenuated. For ΔLat-tubulin (β:F281A mutation), lateral interactions involving the right side of the mutant (viewed from the outside of the microtubule, with plus-end up) are attenuated. Other than the attenuation factors, the mutant subunits are assumed to behave identically as the WT. Attenuation factors were determined by fitting to the mixed tubulin growth rate data.

## ACKNOWLEDGEMENTS

We thank Philipp Kukura for advice on microscope construction and members of the Hancock and Rice labs and Pattipong Wisanpitayakorn for helpful discussions. This work was supported by National Institute of Health grants R01GM076476 to W.O.H. and R01GM098543 to L.M.R, and by National Science Foundation grant MCB-1615938 to L.M.R.. K.J.M was supported by a National Cancer Institute fellowship F99CA223018.

